# Integrated protein-protein interaction and RNA interference screens reveal novel restriction and dependency factors for a tick-borne flavivirus in its human host

**DOI:** 10.1101/2022.11.03.514869

**Authors:** Marion Sourisseau, Yves Unterfinger, Manon Lemasson, Maxime Chazal, Ségolène Gracias, Gregory Caignard, François Piumi, Axel Grot, Sara Moutailler, Damien Vitour, Nolwenn Jouvenet, Muriel Coulpier, Sandrine A. Lacour, Jennifer Richardson

## Abstract

In Europe, tick-borne encephalitis virus (TBEV) is responsible for severe neurological disease in humans. Like other viruses, TBEV is an obligate intracellular life form whose survival requires subversion of metabolic processes and evasion of anti-viral pathways. This feat is achieved in no small part by binary interactions between dedicated viral proteins and host proteins. Such protein-protein interactions (PPI) constitute molecular determinants of critical pathobiologic traits of viruses, including host-range, zoonotic potential and virulence, and represent realistic targets for anti-viral therapies.

To shed light on the pathobiology of TBEV in human, we have resolved the network of PPI established with its human host by interaction proteomics. A high-throughput screen for virus-host PPI was performed involving the complete set of open reading frames of TBEV and the cDNA libraries of *Homo sapiens*, by means of yeast two-hybrid methodology. We have discovered a large set of virus-host protein-protein interactions concerning 42 different human proteins directly interacting with nine viral proteins. Many of these human interactors have never been linked in the literature to viral infection.

The functional significance of the host interactors in viral infection as viral dependency or restriction factors was then characterized *in vitro* by RNA interference, and their function inferred by bioinformatic analysis. Approximately 40% of the identified human proteins have a significative impact on TBEV viral replication. These are engaged in many biological processes, whose involvement in viral infection is expected for many, but enigmatic for some. Further work will be necessary to gain molecular understanding of how these biological processes support or restrict TBEV replication, and whether they constitute viral vulnerabilities that can be exploited therapeutically.

## INTRODUCTION

Flaviviruses, which are principally transmitted by mosquitoes and ticks, cause devastating human diseases such as Dengue fever in the tropics and subtropics and are a constant source of emerging pathogens of pandemic potential, for which ZIKA and Usutu viruses may serve as recent examples. In Europe, a tick-borne flavivirus – the Tick-borne encephalitis virus (TBEV) – is the most significant arbovirus from a medical perspective. Some 3000 cases of TBEV infection, frequently associated with severe central nervous system disease in humans, are reported in Europe each year. Moreover, TBEV is an emerging pathogen on the European continent, as the endemic zone has expanded and the reported incidence increased in many endemic countries^1^. Transmission of TBEV is principally transmitted by the major European tick *Ixodes ricinus*, but may also be food-borne, as TBEV may be acquired by consumption of raw milk and its derivatives from infected sheep and goats in areas where TBE is endemic. Though effective vaccines exist for TBE, they are insufficiently deployed, and antiviral therapy is currently unavailable.

Like all viruses, flaviviruses are obligate intracellular life forms whose survival requires subversion of metabolic circuits and evasion of anti-viral pathways. Within the cell, these pathways are embedded in extensive protein-protein interaction (PPI) networks^2^, and their misappropriation by viruses is largely mediated by binary interactions between dedicated viral proteins and critical network proteins. Indeed, the host proteins targeted by viral proteins tend to occupy positions in the PINs with particular topological features, such that the functional impact of a single interaction may be felt well beyond its immediate neighbourhood by transmission along the cascade of interactions within the PIN^3^. Virus-host PPI thus represent molecular determinants of critical pathobiologic traits of flaviviruses, including host-range, zoonotic potential and virulence. Such interactions represent realistic targets for anti-viral therapies, thus providing a compelling reason to resolve the complete set of virus-cell interactions at the molecular level.

For mosquito-borne flaviviruses of medical significance, virus-host interactions and PPIs in particular have come under intensive study, principally by means of yeast two-hybrid methodology (Y2H) or affinity purification coupled with mass spectrometry (AP-MS)^4,5^. Translation of the flaviviral RNA genome yields a single polyprotein precursor that is cleaved by viral and cellular proteases into three structural (Capsid, prM and Env) and seven non-structural (NS1, NS2A, NS2B, NS3, NS4A, NS4B and NS5) proteins. These have been used in several high-throughput studies to explore PPIs in Y2H^6,7^ or AP-MS^8,9^ screens. The PPIs established between viral and cellular proteins represent molecular effectors of an evolutionary stalemate, certain interactions being conducive while others deleterious to viral survival, and thus representing host “dependency” or “restriction” factors, respectively. Moreover, mapping of PPIs has served to reveal putative molecular determinants of distinct pathobiologic features, such as induction of microcephaly by Zika virus (ZIKV)^8^, as well as druggable targets^9^.

Thus far, the molecular mechanisms that underlie pathobiologic traits have been far less studied in tick-borne flaviviruses than in their mosquito-borne counterparts. A single published study describes a Y2H screen involving the NS3 and NS5 proteins of TBEV and human cDNA libraries^7^. The putative interactions were functionally annotated using *in silico* tools, but their validity was not examined at the biochemical level and their functional implication, such as for viral replication, was not addressed.

In the present study we have performed a high throughput Y2H screen for PPI involving the entire set of open reading frames for the most medically significant arbovirus in Europe, TBEV, and a cDNA library derived from its human host. A set of TBEV-human host PPIs were discovered, many of which having never been previously described for other virus-host systems. The functional significance of the human interactors in viral infection was systematically investigated by performing an RNA interference screen. Many of these interactors were discovered to represent viral restriction or dependency factors – most of which having never been assigned a functional role in other viral infections. The significance of these interactions for the pathobiology of TBEV, in relation to the presumed functional role of the human interactors, is discussed. Our study thus illuminates the strategies by which tick-borne flaviviruses control cellular processes and cause disease, and should ultimately contribute to disclosing viral vulnerabilities that can be exploited therapeutically.

## RESULTS

### The TBEV – *Homo sapiens* protein-protein interaction network

In order to resolve the PPI network between TBEV and its human host, a *H. sapiens* cDNA library was screened by Y2H assay for binary PPI with the complete set of ORFs of TBEV. These encoded the three major structural – C, prM, and E – and eight nonstructural proteins – NS1, NS2A, NS2B, NS3, NS4A, 2K, NS4B and NS5 (Fig. 1a). ORFs encoding the cleavage products of full-length prM, namely, the mature M protein and the cleaved N-terminal portion (prM’) of prM, were used separately for screening. Among 231 yeast colonies selected for sequencing, 27 human proteins interacting with TBEV bait proteins were identified. Of note, the screen described in the current report was performed concomitantly with a Y2H screen for PPI between the *H. sapiens* proteome and the complete set of ORFs of the Louping Ill virus (LIV), a tick-borne flavivirus closely related to TBEV and an important pathogen in sheep, as well as with Y2H screens for PPI between the *Bos taurus* proteome and the complete set of ORFs of both TBEV and LIV (data not shown). Based on PPI disclosed between LIV bait proteins and human prey or between TBEV bait proteins and bovine preys in these independent Y2H screens, orthologous PPI between TBEV bait proteins and certain human proteins were specifically evaluated by Gap repair (GR) assay (Fig. 1b).

**Fig.1.**
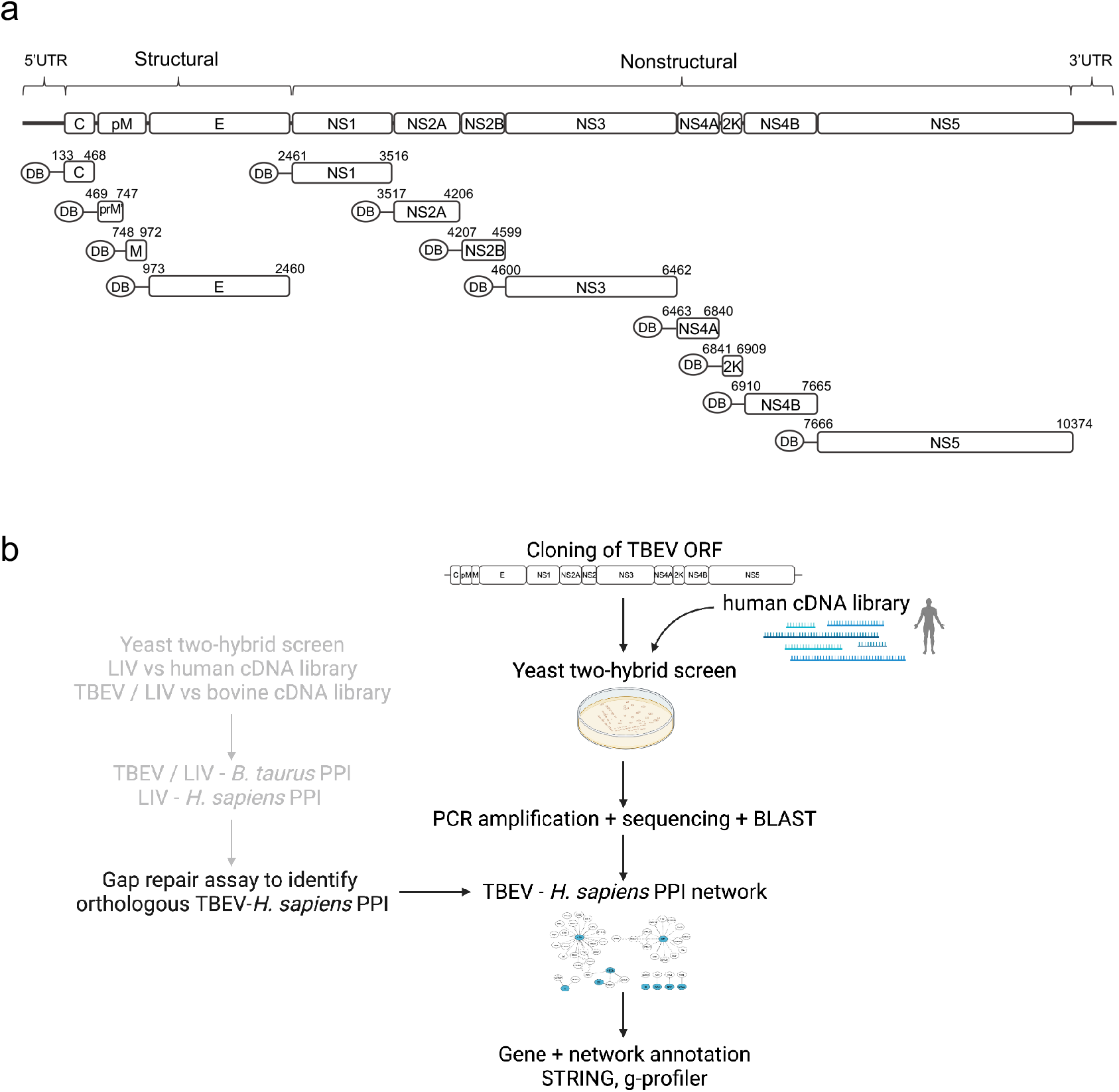
Yeast two-hybrid (Y2H) workflow for discovery of TBEV-human host protein-protein interactions. **a**, Schematic representation of the TBEV RNA genome depicting viral ORFs cloned downstream of the DNA-binding domain (DB) of the Gal4 transcription factor, for expression of viral baits in the Y2H screen. b, Experimental workflow for Y2H screens and gap repair assays.

PPI observed in the original TBEV - H. sapiens screen supported by retrieval of a single sequence were also evaluated by GR. This strategy let to confirmation of nine human proteins originally identified in the TBEV-human host Y2H screen as interacting with TBEV bait proteins, and to identification of fifteen additional human proteins that interact with TBEV bait proteins (Supplementary Table 1). The resulting set of 42 human prey proteins are shown in a network representation in which nodes, representing proteins, are connected by edges, representing binary PPI (Fig. 2a). Nine nodes represent viral bait proteins, while 42 represent human preys. Among these last, 20 interacted with NS5, 13 with prM’, 3 with NS2B, 2 with C and one each with M, NS1, NS3 and NS4A and 2K (Fig. 2b). One human prey protein, encoded by the UBQLN1 gene, interacted with both 2K and NS2B proteins. PPI were not evidenced for E, NS2A and NS4B proteins.

**Fig.2.**
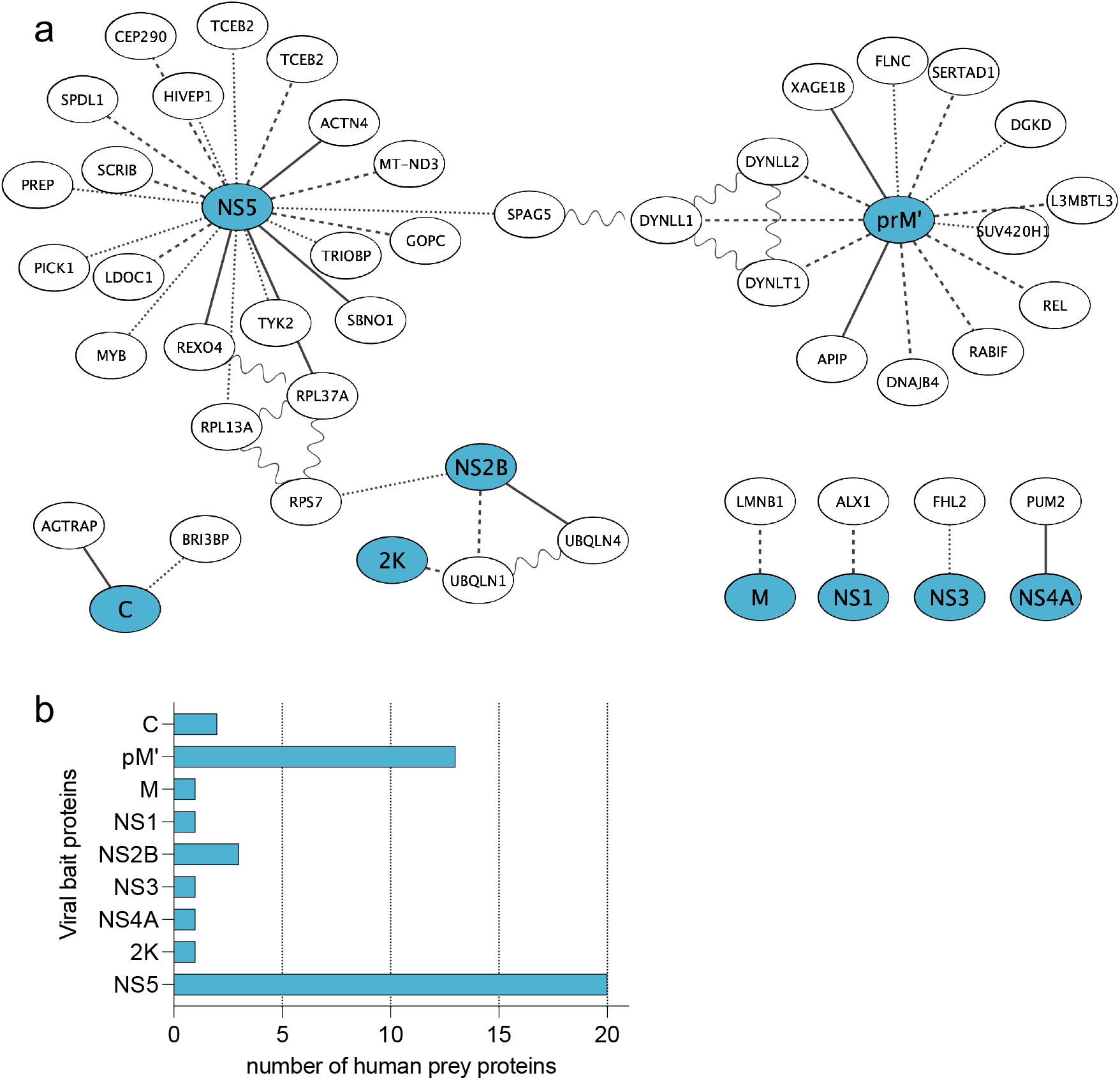
TBEV-human host protein-protein interactions revealed by Y2H. **a**, TBEV-human host protein-protein interaction (PPI) network. Nodes and edges represent proteins (viral bait and human prey) and PPI, respectively. Regarding edges, dots, equal dashes and solid line types represent PPI identified by gap repair (GR) only, by Y2H only or by both GR and Y2H, respectively. Sine waves represent physical PPI between human prey proteins retrieved from the STRING database^12^. **b**, Number of distinct *H. sapiens* prey proteins identified for each viral bait protein.

### Overlap of PPI with published TBEV-*H. sapiens* PPI

In a previous study, a Y2H screen had been carried out between the NS3 and NS5 proteins of TBEV (strain 263^10^) and human cDNA libraries^7^. Full-length NS5, as well as domains thereof responsible for RNA polymerase and methyltransferase functions, were used separately for screening. The set of human proteins interacting with NS5 (NS5 bait set) from Le Breton *et al*.^7^ and the present study were merged (Fig. 3a). Of the 33 and 20 human prey revealed in the former and present study for NS5, respectively, only 3 – encoded by GOPC, SCRIB and TYK2 genes – were common to both studies.

**Fig.3.**
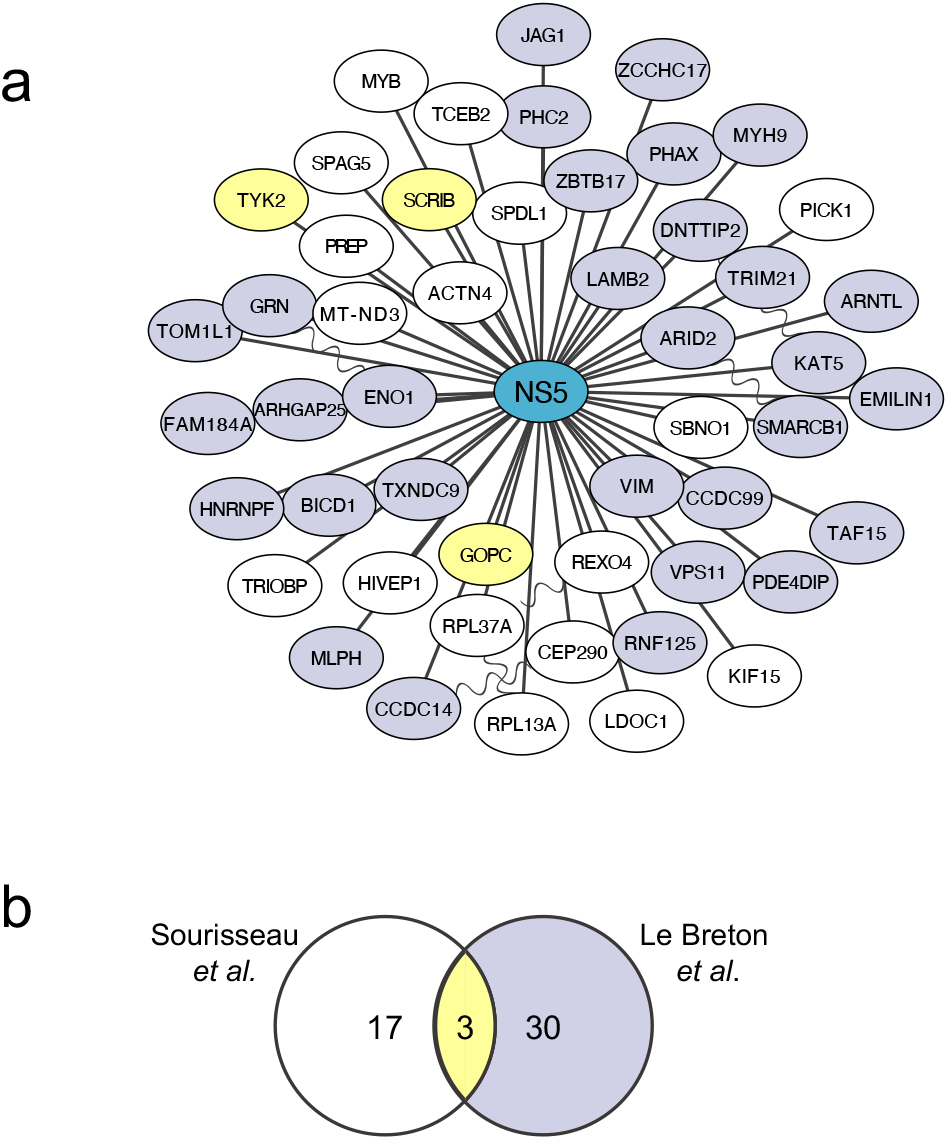
Comparison of TBEV NS5-human host protein-protein interactions uncovered in independent Y2H screens. **a**, TBEV NS5-human host PPI network including data from both Le Breton *et al*.^7^ and the present study. Nodes and edges represent proteins (TBEV NS5 and human prey) and PPI, respectively. Nodes with purple or white fill colour represent prey proteins found only by Le Breton *at al*.^7^ or only in the present study, respectively, while nodes with yellow fill colour represent prey proteins found in both studies. Sine waves represent physical PPI between human prey proteins retrieved from STRING^12^. **b**, Venn diagram showing overlap (yellow) in human prey proteins identified in Le Breton *et al*.^7^ (purple) and in the present study (white).

### Biological processes engaged by TBEV infection

To identify the biological processes engaged by TBEV infection, g:Profiler^11^ was used to retrieve enriched Gene ontology terms for the human prey proteins disclosed in the present study. Enrichment analyses of the prey proteins, collectively or by individual bait set, retrieved few or no enriched GO terms, respectively, that attained statistical significance. Thus, to identify major biological processes engaged by TBEV infection, the same analyses were performed for individual bait sets after extension of each set by the addition of up to ten human proteins (Supplementary Table 2) that physically interact with each prey in the bait set (Fig. 4b), as retrieved from STRING^12^.

**Fig.4.**
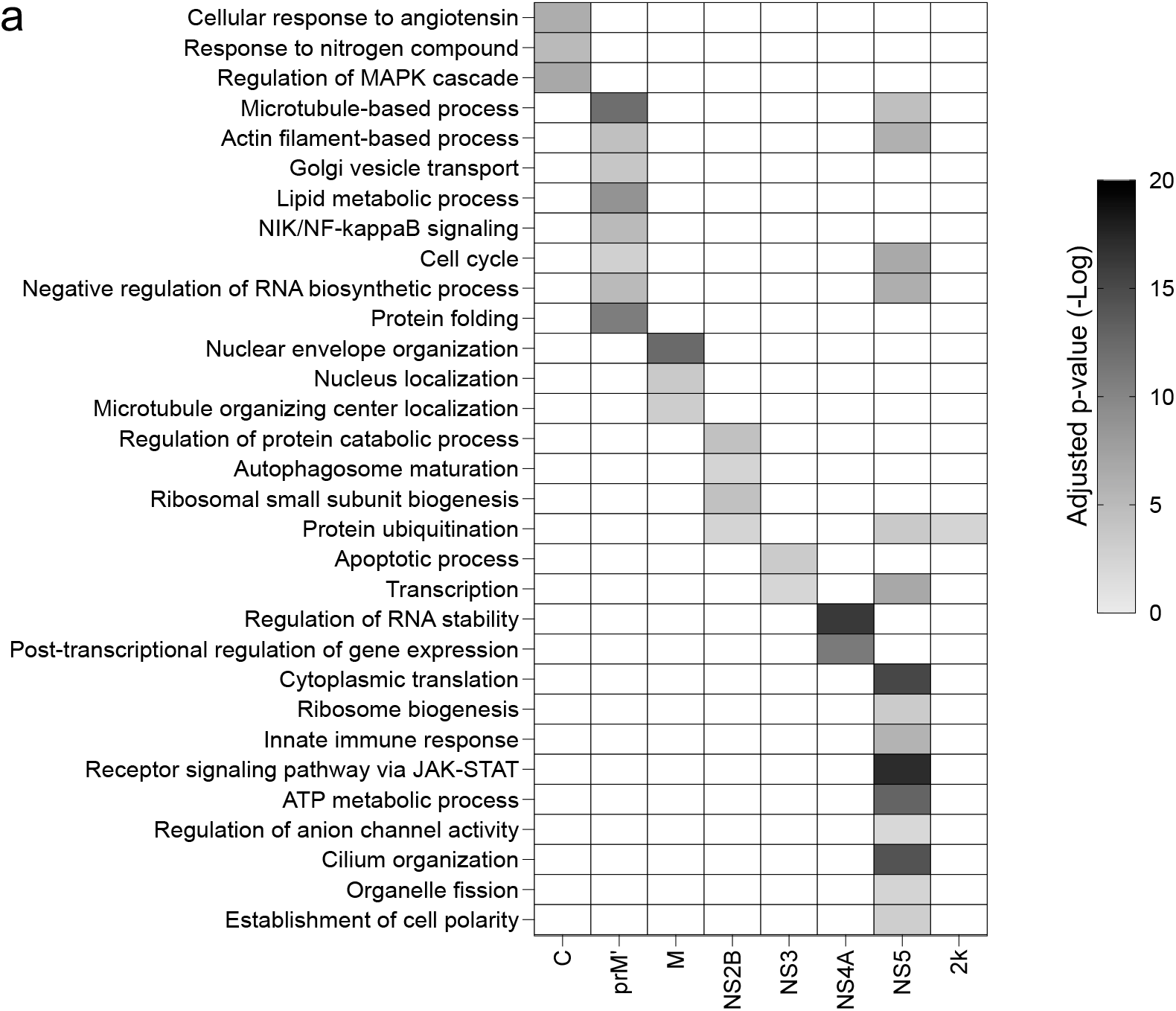
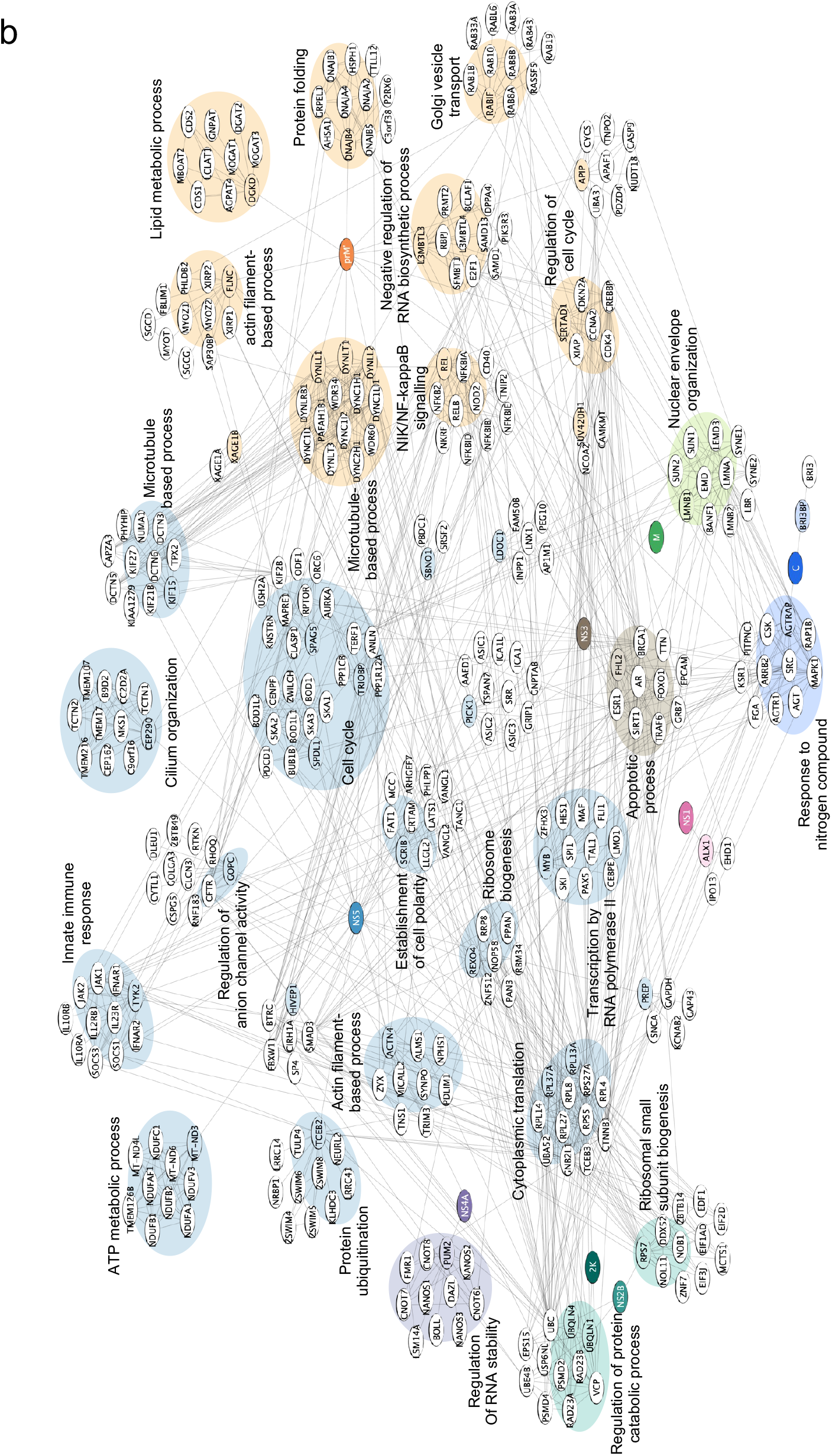
Functional annotation of extended TBEV-human host protein-protein interaction network. Extended TBEV-human host PPI network displaying Gene Ontology annotations. For extension of the network of PPI disclosed in the present study, PPI of human prey proteins, up to 10 for each prey of each bait set, were retrieved from the STRING database^12^ and merged with the original PPI network deduced from Y2H and GR assay (present study only). Gene ontology terms were retrieved for the extended version of individual bait sets using g:Profiler^11^. a, A subset of the most statistically significant GO-Biological pathways, along with their associated p-value, is displayed for each extended bait set. b, Network representation and associated GO terms after extension. Each bait set of prey proteins is distinguished by functional “clouds” of a unique colour. Nodes representing human prey proteins disclosed by the Y2H and GR screen are filled, while nodes representing proteins added by String extension are unfilled.

Multiple enriched GO terms that attained statistical significance were retrieved for the STRING-extended bait sets (Supplementary Table 3). Those concerning the most salient biological processes are shown for individual bait sets in Fig. 4a and illustrated in network format in Fig. 4b. Among them, we identified certain cellular processes with evident links to viral infection, such as cytoskeleton component-based processes (NS5, prM’ and M), vesicle transport (C and prM’), translation (NS2B, NS5), apoptotic processes (NS3), protein degradation (NS2B) and innate immune response (NS5) (Fig. 4 and Supplementary Table 3). Of note, enriched GO terms displayed little overlap between extended bait sets, suggesting that the biological processes engaged by individual viral proteins are largely distinct.

### Functional analysis of TBEV-human host PPI

To assess the role of the human prey proteins identified in the present study in the TBEV life cycle, we analyzed the consequence for viral replication of silencing their expression by RNA interference (RNAi). Small interfering RNA (siRNA) was used to transfect A549 cells prior to infection with TBEV. Viral replication was assessed 24h after infection by quantification of infectious particles released into the supernatant (Fig. 5a). Transfection with irrelevant (non-targeting) siRNA (siNT) was used to provide a reference value. Infectious titers measured after silencing with siRNA specific for individual human prey proteins were expressed relative to titers measured after treatment with siNT, and plotted against their respective p-value. When a specific siRNA induced at least a two-fold change in normalized infectivity with a p-value <0.01, the targeted host factor was considered a hit. Host factors whose silencing diminished titers were considered to represent dependency factors, while those whose silencing augmented titers were considered to represent restriction factors.

**Fig.5.**
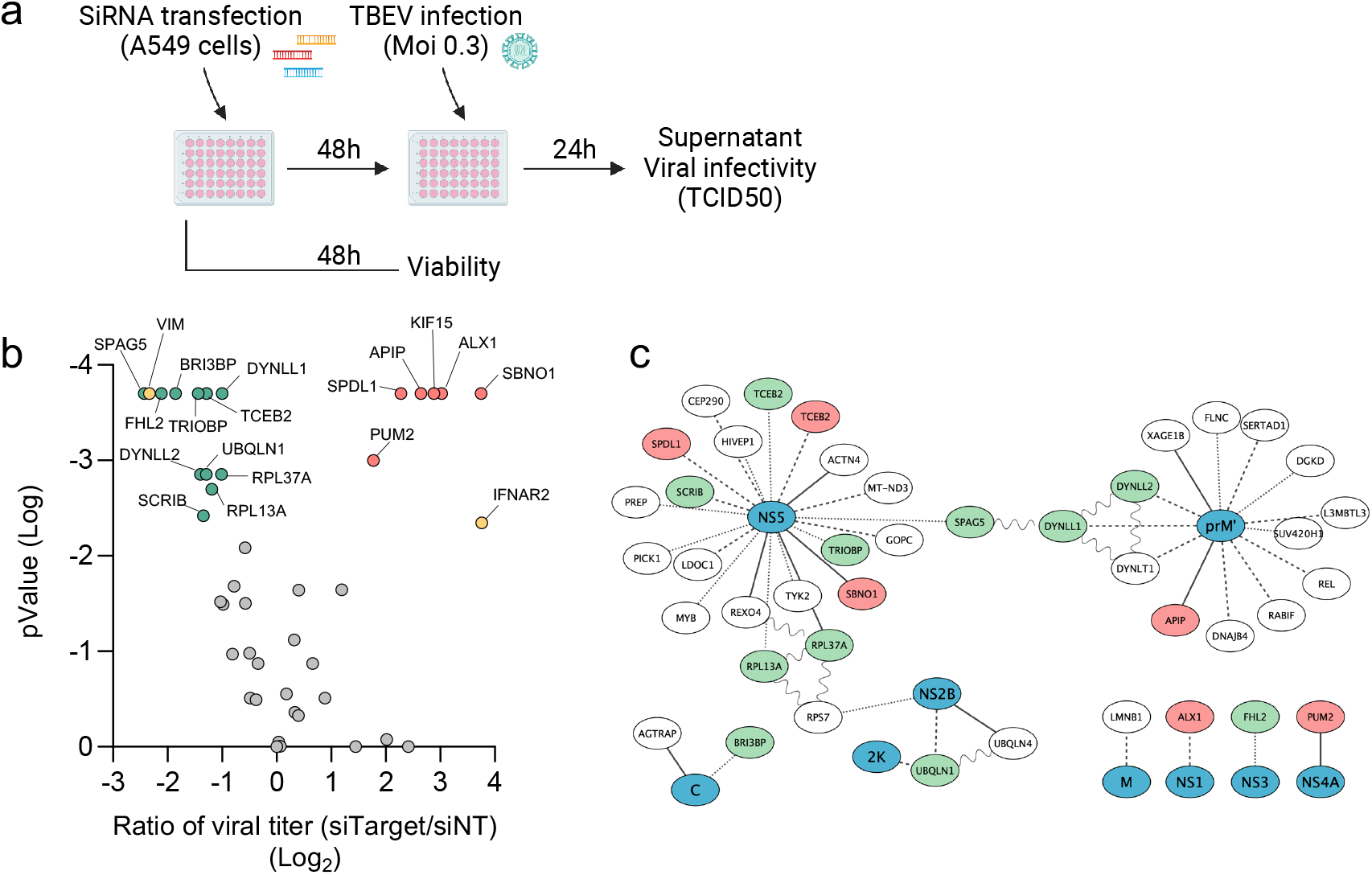
RNAi screen to identify TBEV host dependency and restriction factors. **a**, Experimental workflow for the loss-of-function RNAi screen. **b**, A549 cells were transfected with an siRNA library and challenged with TBEV (MOI 0.5). Infectivity titers in the supernatant, 24h post-infection, were quantified by end-point dilution and normalized to infectivity titers obtained for cells transfected with nontargeted siRNA (Log_2_ fold change). Data, expressed as TCID_50_, are means of 2 biological replicates (with technical replicates). P-values were calculated by comparison of data obtained from cells transfected with targeted and nontargeted siRNA (Mann-Whitney test). Host dependency and host restriction factors revealed in the present study are marked in green and red, respectively, while positive controls for dependency and restriction factors (VIM and IFNAR2) are shown in yellow. **c**, TBEV-human host PPI network showing host dependency factors in green and host restriction factors in red.

As expected, siRNA directed against the restriction and dependency control proteins IFNAR2 and VIM^13^ modulated infection accordingly. Small interfering RNA against IFNAR2 induced a 13-fold increase in viral release, while VIM extinction significantly decreased infectivity titers by a 5.3-fold factor. Among the 42 host proteins tested (Supplementary Table 4), the silencing of 19 of them significantly modulated infectious particle release by 2-fold or greater, as compared with the reference titer (Fig. 5b).

Eleven hits for which silencing negatively affected TBEV release (encoded by SPAG5, FHL2, BRI3BP, TCEB2, TRIOBP, DYNLL1, DYNLL2, UBQLN1, RPL37A, RPL13A and SCRIB genes) may be considered to represent TBEV dependency factors. Six factors, for which silencing enhanced TBEV release by more than 2-fold, qualified as restriction factors (encoded by PUM2, SPDL1, APIP, KIF15, ALX1 and SBNO1 genes).

## DISCUSSION

In comparison with their mosquito-borne counterparts, tick-borne flaviviruses are notoriously understudied, including as regards the PPI established between viral and host proteins, which permit viruses to co-opt biological circuits and circumvent the host’s antiviral response. In the present study we have performed a Y2H screen between the most medically important arbovirus in Europe, TBEV, and its human host, thereby revealing multiple hitherto unknown PPI. The functional role of the human interactors (“prey proteins”) was systematically addressed by RNA interference, disclosing multiple viral restriction and dependency factors, many of which undescribed for other viruses.

Among the 42 human prey proteins of TBEV revealed in the present Y2H screen and associated gap repair analyses, 6 (14%) and 11 (26%) were found to be restriction and dependency factors, respectively. Thus, a high proportion (≈ 40%) of human proteins that physically interact with TBEV proteins could be shown to have a measurable impact on viral replication. This percentage is likely to be underestimated, as knockdown of certain proteins may have been inadequate, due for example to high levels of mRNA or to an extended half-life of the protein product. Moreover, knockdown of certain proteins may have been compensated by non-targeted human proteins playing a redundant role in viral replication.

Of the three structural proteins of TBEV that were used for screening, C, prM (as its cleaved products prM’ and M) and E, human prey proteins were discovered for C, prM’ and M. Following expression of the precursor polyprotein of flaviviruses in the ER membrane and cleavage by the viral protease, the C protein persists as an ER membrane-anchored protein facing into the cytosol, until further proteolytic cleavage releases soluble C into the cytosol. The mature form of flaviviral C may strongly associate with the ER-derived membranes at the site of viral replication^14^, but can also be imported into the nucleus^15^. Both of the human prey proteins found for TBEV C protein – Angiotensin II Receptor Associated Protein or AGTRAP and BRI3 (Brain protein I3) Binding Protein (BRI3BP) – are multi-pass membrane proteins. The latter was found to be a dependency factor. To the best of our knowledge, neither protein has previously been found to bind to viral proteins or to play a role in viral replication. BRI3BP, presumed to be principally expressed in the mitochondrial membrane, is virtually uncharacterized, though one study suggests that it might have pro-apoptotic activity^16^.

Immature flaviviral particles bud into the lumen of the ER and follow the exocytosis pathway for ultimate egress via the plasma membrane. During viral trafficking the prM protein of flaviviruses is expressed as a membrane-anchored protein, until cleavage by the cellular endoprotease furin in the trans-golgi network releases the NH2-terminal portion, prM’, into the ER lumen, leaving the mature M protein anchored in the viral membrane. Twelve prey proteins were evidenced for the prM’ moiety; namely, Apoptotic peptidase activating factor 1 interacting protein (APIP), Diacylglycerol kinase delta (DGKD), DnaJ heat shock protein family (Hsp40) Member B4 (DNAJB4), dynein light chain 1, cytoplasmic (DYNLL1), Dynein light chain 2, cytoplasmic (DYNLL2), Dynein light chain Tctex-type 1 (DYNLT1), Filamin C (FLNC), Histone methyl-lysine binding protein 3 (L3MBTL), RAB Interacting Factor (RABIF), c-Rel (REL), SERTA domain-containing protein 1 (SERTAD1), Suppressor of variegation 4-20 homolog 1 (SUV420H1) and X antigen family member 1 (XAGE1B). Among these twelve human prey proteins, APIP was revealed to be a viral restriction factor, while the paralogs DYNLL1 and DYNLL2 were evidenced as dependency factors. To the best of our knowledge, APIP has not previously been reported to bind to viral proteins or to play a role in viral replication. APIP is a 5-methylthioribulose-1-phosphate dehydratase belonging to the methionine salvage pathway^17,18^, and an inhibitor of both apoptosis and pyroptosis^17^.

The dynein light chains DYNLL1, DYNLL2 and DYNLT1 are non-catalytic accessory components of the cytoplasmic dynein 1 complex, and presumed to link dynein to various cargos for retrograde motility of vesicles along microtubules. Various proteins have been shown to physically interact with dynein light chains (for review see ^19^). Our previous study disclosed a physical interaction between the prM’ moiety of TBEV and the DYNLL1 ortholog in the tick *Ixodes ricinus*^20^, suggesting that the interaction, and possibly functional role, is conserved across in the arboviral life cycle between mammalian host and arthropod vector.

A member of the NF-κB family of transcription factors, the c-Rel protein is involved in the innate immune response to myriad viruses, and has been shown to physically interact with the Tax protein of human T-cell leukemia virus^21^. The transcriptional factor SERTAD1 has been shown to interact with the E2 protein of Classical swine fever virus^22^, a member of the genus *Pestivirus* in the family *Flaviviridae*, and ablation of the interaction restricts replication in cellular models and *in vivo* in swine^23^. SUV420H1, a dimethyl transferase of histone H4, has been described to be involved in the latency of Epstein barr virus^24^. The remaining human prey proteins of prM’, DGKD, DNAJB4, FLNC, L3MBTL3 and RABIF, have not, to our best knowledge, been implicated as intervening in the life cycle of viruses.

Of the 8 ORFs encoding nonstructural proteins of TBEV that were used for screening, human prey proteins were discovered for NS1, NS2B, NS3, NS4A, 2K and NS5.

ALX homeobox protein 1 (ALX1), the unique interactor of NS1 in our Y2H screen, exerts a negative impact on TBEV infection. This transcription factor has been shown to play a critical role in coordinating the antiviral innate immune response, along with known virus activated factors such as IRF3, IRF7, STAT1, and/or NF-kB^25^.

Three human prey proteins were discovered for the NS2B protein; namely, the ribosomal protein S7 (RPS7) and two members of the ubiquilin family (UBQLN1 and UBQLN4), UBQLN1 being identified as a dependency factor for TBEV. Ubiquilins behave as adaptor proteins that target poly-ubiquinated proteins to the proteosome for degradation, and have also been implicated in autophagy and endoplasmic reticulum-associated protein degradation (reviewed in ^26^). Of note, proteins of several viruses have been found to interact with ubiquilin family members^27,28,29^, in some cases providing benefit for viral replication^29^. In our previous study, the tick *I. ricinus* ortholog of ubiquilin-1 (UBQLN1 gene), was found by Y2H to be an interactor of TBEV NS4A^20^, suggesting that co-opting of ubiquilin by TBEV may be conserved in its mammalian host and arthropod vector.

The NS3 protein is not only a viral protease required for cleavage of the viral polyprotein precursor and of release of individual nonstructural proteins, but also an RNA helicase involved in viral RNA synthesis^30,31^. Four-and-a-half LIM domain protein 2 (FHL2), the sole interactor of NS3 in our Y2H screen, was identified as a dependency factor for TBEV. FHL2 has been shown to be implicated in apoptosis^32,33^, a process shown to be modulated by NS3 protein of multiple flaviviruses^34,35^.

The TBEV NS5 protein plays an essential role in viral replication as both a methyltransferase and an RNA-dependent RNA polymerase. By inhibiting the type-I IFN pathway, it also plays on important role in immune evasion^36–38^. Twenty human prey proteins were evidenced for the TBEV protein (see Fig 3); of which 6 and 3 were identified as dependency and restriction factors, respectively. Dependency factors included the ribosomal proteins L37A and L13A (RPL37A and RPL13A, respectively), Scribble (SCRIB), TRIO And F-Actin Binding Protein (TRIOBP), Elongin B (TCEB2) and Sperm Associated Antigen 5 (SPAG5), while Strawberry Notch 1 (SBNO1), Kinesin Family Member 15 (KIF15) and Spindle Apparatus Coiled-Coil Protein 1 (SPDL1) behaved as restriction factors.

Regarding the dependency factors that bound to NS5, both RPL37A and RPL13A are components of the large 60S subunit, with the latter playing a role in downregulation of the inflammatory gene expression as a component of the IFN-gamma-activated inhibitor of translation (GAIT) complex (reviewed in ^39^). RPL13A silencing also impacts replication of influenza^40^. The physical association between NS5 and SCRIB, previously described in Le Breton *et al*.^7^, has been suggested to antagonize the innate immune response by impeding interferon-stimulated JAK-STAT signalling^41^. The present study, however, is the first to evidence an impact for SCRIB in TBEV infection.

TRIOBP interacts with the protein Trio, implicated in neural tissue development, organization of the actin cytoskeleton, cell motility and cell growth and with F-actin, with no described role in viral infection up to now. TCEB2 is a regulatory subunit of the transcription factor B (SIII) complex, known to be subverted by HIV to co-opt proteosomal degradation pathways to counteract cellular restriction factors^42^

SPAG5, originally identified as a microtubule-associated protein with dual localization to both centrosomes and kinetochores, is required for centrosome integrity, mitotic spindle formation and chromosome segregation during mitosis^43,44^. It also has an independent role in regulating apoptosis induced by cell stress^43,44^. NS5 of Dengue and ZIKA viruses (DENV and ZIKV) are known to localize at centrosomes during cell division^45,46^. While formation of the ZIKV viroplasm is closely associated with the centrosome and the Golgi MTOC^47^, neither viroplasm formation nor virus production are significantly impaired in the absence of centrosomes. Thus, the functional significance of interactions between flavivirus proteins with centrosomes remains ill-defined.

Regarding the dependency factors that bound to NS5, SBNO1 has been shown to associate with nonstructural viral proteins from several RNA+ viruses, such as ZIKV NS4A and NS5^8,48^, as well as Sars-Cov2 Nsp12^49^. Orthologs of SBNO1 are implicated in nervous system development involving the Notch pathway, or in activation of Notch signaling pathways in zebrafish^50,51^ and *Drosophila*^53–55^, respectively. ZIKV interferes with neuronal differentiation by dysregulating the canonical Notch signalling pathway, both in cultures of hNPCs and in mouse embryonic brains^52^. An impact for SBNO1 on viral replication, however, had never previously been reported. The two remaining NS5-binding restriction factors, KIF15 and SPDL1, both located in kinetochores, have not previously been linked to viral infection.

In conclusion, our study has disclosed novel PPIs for a highly significant European arbovirus and revealed multiple viral restriction and dependency factors, some shared with other viruses while others never previously described in the published literature. Beyond elucidation of the strategies by which tick-borne flaviviruses control cellular processes and cause disease, our study sheds light on the cellular function of certain host interactors whose biological role is as yet enigmatic.

## ACKNOWLEDGEMENTS

This study was made possible by a grant from the French national research agency (Project N° ANR-19-CE35-0015-01). The Ph.D. of ML was financed by a grant from the French Agency for Food, Environmental and Occupational Health & Safety (ANSES, Grant No. PHD2017-2020). MS received post-doctoral funding from the DIM1Health. The Ph.D. of AG is financed by the LabEx *Integrative Biology of Emerging Infectious Diseases*.

## AUTHORS’ CONTRIBUTIONS

JR, SAL, MuC, NJ and MS conceived and designed the study. MS, YU, MaC, SG, ML and AG carried out the wet lab experiments and/or bioinformatic analyses. GC and DV provided guidance and assistance with the Y2H screen. FP provided support for bioinformatic analyses. SM provided an essential resource. The paper was drafted by JR and MS and revised by DV, GC, ML, MuC and SM. All authors read and approved the final manuscript.

## MATERIALS & METHODS

### Cell culture

The simian epithelial cell line Vero (ATCC No. CCL-81), used for propagation and titration of TBEV, and the human epithelial cell line A549 (ATCC No. CCL-185) were cultivated in Dulbecco’s modified Eagle’s medium (DMEM; Gibco BRL Life Technologies, Gaithersburg, MD) supplemented with 5% fetal calf serum (FCS) (Eurobio) and 100 U/ml penicillin and 100 μg/ml streptomycin (both from Gibco BRL Life Technologies). Cells were maintained at 37°C in the presence of 5% CO_2_.

### Viral propagation and quantification

The Hypr strain of TBEV (GenBank accession number U39292.1), isolated in the Czech Republic from a child with TBE^53^, was propagated and titrated in Vero cells grown in DMEM supplemented with antibiotics as described above and 2% FCS. Viral stocks were prepared as described in Lemasson *et al*.^20^ and quantified by endpoint dilution assay on Vero cells, as visualized by the virus-induced cytopathic effect (CPE), as previously described^54^. Briefly, 1.5 × 10^4^ Vero cells were seeded in 100 μL of DMEM supplemented with antibiotics and 2% FCS in wells of a 96-well plate the day prior to infection with 25 μL of serial dilutions of virus in medium as above (6 wells per dilution). Five days post-infection, infected wells were scored for evidence of virus-induced CPE. The fifty percent tissue culture dose (TCID_50_) was calculated according to the method of Reed and Muench^55^.

### Yeast two-hybrid screen

In order to identify human proteins that physically interact with TBEV proteins, a yeast two-hybrid (Y2H) screen^56^ was performed. To this end, the coding sequences of all open reading frames of TBEV Hypr were cloned and recombined into the pPC97 plasmid (Invitrogen) downstream of the DNA-binding domain of the Gal4 transcription factor (Gal4-DB; Fig. 1), as described in Lemasson *et al*.^20^. Of note, ORFs encoding the cleavage products of full-length prM, namely, the mature M protein and the cleaved N-terminal portion (prM’) of prM, rather than the entire prM protein, were used separately for screening. A human cDNA library, in which cDNA derived from A549 cells had been cloned within the pDEST22 plasmid (Invitrogen) for in frame expression downstream of the activation domain of Gal4 (Gal4-AD), was kindly provided by Pierre-Olivier Vidalain. After transformation of the Y2H Gold and Y187 yeast strains with pPC97 constructs encoding viral baits and the cDNA library encoding *H. sapiens* preys, respectively, the Y2H screen was conducted essentially as described by Lemasson *et al*^20^. The TBEV-*H. sapiens* screen was part of a broader screen in which viral bait proteins from not only TBEV but also from the highly related Louping Ill virus (LIV) were used to screen cDNA libraries from not only *H. sapiens*, but also *Bos taurus* species. The cloning of LIV baits and the derivation of the *B. taurus* cDNA library have been described in Lemasson *et al*.^20^ and Fablet *et al*.^57^, respectively.

### Gap Repair

To determine whether PPI identified for proteins of LIV but not TBEV in Y2H screens of the human cDNA library were actually shared by the two viruses, interaction between the orthologous TBEV proteins and LIV-binding prey was evaluated by gap repair (GR) assay^58^ as described in Lemasson *et al*^20^. Briefly, yeast carrying plasmids expressing DB-fused viral proteins were co-transformed with 10 ng of linearized pPC86 plasmid and 3 μL of PCR product encoding the protein of interest (as produced during the Y2H screen) and plated on medium lacking leucine (leu) and tryptophan (trp) and then transferred to medium differing by the absence of histidine (his) and the presence of 5 mM 3-amino-1,2,4-triazole (3-AT; Sigma-Aldrich, Saint-Louis, MO, USA). Homologous recombination between pPC86 and the PCR product then reconstituted the expression cassette encoding the AD-fused prey protein, and growth on medium lacking leu, trp and his was conditioned by physical interaction between viral and *H. sapiens* proteins. A similar strategy was used to determine whether PPI identified in Y2H screens for LIV or TBEV with the bovine but not human cDNA library were actually shared by the two mammalian species. In brief, based on sequence analysis of the PCR products corresponding to bovine prey, the orthologous regions of human cDNA were amplified from mRNA of A549 cells by RT-PCR. These PCR products were then used in GR assay as described above.

### RNAi Screen

RNAi targeting the 42 human proteins (a pool of 4 siRNA for each target) recovered in the Y2H assay (RNAi sequences provided in Supplementary Table 4) was purchased from Horizon Discovery (Cherry-pick Library in SMARTpool format). Three days prior to transfection, 10^4^ A549 cells were seeded in wells of 48-well plates. Transfection was performed in triplicate with 25 nM of siRNA and 1 μL of DharmaFECT 1 Transfection reagent (Horizon Discovery) as per manufacturer’s instructions. Forty-eight hours post-transfection, RNAi-transfected cells were infected with TBEV Hypr at an MOI of 0.5 TCID_50_ per cell in DMEM supplemented with antibiotics 2% FCS. Viral inoculum was removed 90 minutes later, and cells were washed twice with PBS before addition of 200 μL of medium as above. Supernatants were collected 24h post-infection. The impact of RNAi on viral infection was assessed by quantification of infectious particles in supernatants of A549 cells by endpoint dilution assay on Vero cells, and expressed in TCID_50_ as the mean of biological triplicates.

Cell viability after knockdown was evaluated for each siRNA pool in the absence of infection, 48h post-transfection of A549, by using commercially available reagents (Cell Titer Glo Luminescent Cell Viability Assay; Promega) according to the manufacturer’s instructions (data not shown).

### In silico and Gene Ontology (GO) analyses

PPI were graphically represented using Cytoscape software^59^. The g:GOSt tool of g:Profiler ^11^(https://biit.cs.ut.ee/gprofiler/gost) was used for functional enrichment analysis of our bait-sets, using g:SCS algorithm with a defined p-value threshold of 0.01. Human prey proteins were submitted to the Gene Ontology (GO) term enrichment service using an *H. sapiens* organism filter to identify biological processes enriched in the yeast two-hybrid dataset. As few statistically significant terms were recovered, sets of human prey proteins for individual viral bait proteins (“bait sets”) were extended by retrieval of up to 10 physical PPI for each prey of each bait set from the STRING database^12^, using the Cytoscape StringApp^60^. The list of GO terms and associated p-values was condensed by using the REVIGO (reduce and visualize Gene Ontology; http://revigo.irb.hr/)^61^.

### Statistical analyses

Data from the siRNA screen were analyzed for statistical significance with Prism9 software (GraphPad Software) using the Mann-Whitney test. A p-value < 0.01 was considered significant. Values represent the mean and standard deviation (SD) of at least two independent experiments, each performed in biological triplicate.

Enriched GO terms were obtained using the g:SCS (Set Counts and Sizes) correction method. Adjusted p-values < 0.01 were considered significant.

## SUPPLEMENTARY DATA

**Supplementary Table 1**:Summary of TBEV-*H. sapiens* protein-protein interactions (PPI) identified by yeast two-hybrid (Y2H) screening. Columns A and B display TBEV bait proteins and cognate human prey proteins, respectively, the latter referring to STRING identifiers. Uniprot identifiers and Uniprot protein descriptions are provided in columns C and D, respectively. Identification (+) or not (-) of the PPI in the Y2H screen is indicated in column E, and the number of yeast clones recovered for each PPI is indicated in column F. PPI that were evidenced by Gap repair (GR) assay, evaluated either because few yeast clones had been selected or because the PPI had been identified for LIV and the prey protein in question, are indicated in column G. Similarly, some PPI were evidenced by GR assay subsequent to their identification in a screen for PPI between TBEV and the bovine host, as indicated in column H. A summary of the evidence for each PPI (Y2H screen or GR assay or both) is provided in column I.

**Supplementary Table 2**:Bait sets and Extended bait sets. In order to return statistically significant Gene Ontology terms, bait sets for individual viral proteins were extended by the addition of up to ten PPI for each human prey of each bait set, as retrieved from the STRING database^12^, using the Cytoscape StringApp^60^.

**Supplementary Table 3**:Enriched Gene ontology terms for extended viral bait sets. Enriched gene ontology terms as regards Biological processes were retrieved for the STRING-extended version of individual using g:Profiler^11^.

**Supplementary Table 4**:Sequence of siRNA used to knock down expression of genes encoding human proteins that physically interact with TBEV proteins, as determined in the Y2H screen described in the present study.

